# Pathogen Reduction of SARS-CoV-2 Virus in Plasma and Whole Blood using Riboflavin and UV Light

**DOI:** 10.1101/2020.05.03.074971

**Authors:** Izabela Ragan, Lindsay Hartson, Heather Pidcoke, Richard Bowen, Raymond P. Goodrich

**Affiliations:** Department of Biomedical Sciences, Colorado State University, Fort Collins, Colorado 80523; Infectious Disease Research Center, Colorado State University, Fort Collins, Colorado 80523; Translational Medicine Institute, Colorado State University, Fort Collins, Colorado 80523

## Abstract

**BACKGROUND:** Severe Acute Respiratory Syndrome Coronavirus 2 (SARS-CoV-2) has recently been identified as the causative agent for Coronavirus Disease 2019 (COVID-19). The ability of this agent to be transmitted by blood transfusion has not been documented, although viral RNA has been detected in serum. Exposure to treatment with riboflavin and ultraviolet light (R + UV) reduces blood-borne pathogens while maintaining blood product quality. Here, we report on the efficacy of R + UV in reducing SARS-CoV-2 infectivity when tested in human plasma and whole blood products.

**STUDY DESIGN AND METHODS:** SARS-CoV-2 (isolate USA-WA1/2020) was used to inoculate plasma and whole blood units that then underwent treatment with riboflavin and UV light (Mirasol Pathogen Reduction Technology System, Terumo BCT, Lakewood, CO). The infectious titers of SARS-CoV-2 in the samples before and after R + UV treatment were determined by plaque assay on Vero cells. Each plasma pool (n=9) underwent R + UV treatment performed in triplicate using individual units of plasma and then repeated using individual whole blood donations (n=3).

**RESULTS:** Riboflavin and UV light reduced the infectious titer of SARS-CoV-2 below the limit of detection for plasma products at 60-100% of the recommended energy dose. At the UV light dose recommended by the manufacturer, the mean log reductions in the viral titers were ≥ 4.79 ± 0.15 Logs in plasma and 3.30 ± 0.26 in whole blood units.

**CONCLUSION:** Riboflavin and UV light effectively reduced the titer of SARS-CoV-2 to the limit of detection in human plasma and by 3.30 ± 0.26 on average in whole blood. Two clades of SARS-CoV-2 have been described and questions remain about whether exposure to one strain confers strong immunity to the other. Pathogen-reduced blood products may be a safer option for critically ill patients with COVID-19, particularly those in high-risk categories.

## INTRODUCTION

Due to a combination of factors including rapidly mutating viral strains, increasingly close animal-human contacts, and ever burgeoning rates of travel and urbanization, the rate at which new human pathogens emerge and spread globally appears to be rising over the last 80 years (1). Climate change is likely to accelerate pandemic emergence because temperate zones encompass a larger area of the globe, expanding vector territories and favoring bacterial pathogens (2,3). Mass gatherings and higher Basic Reproduction Numbers further contribute to rapid dissemination around the globe (4,5). The emergence of Coronavirus Disease 2019 (COVID-19), whose causative agent is Severe Acute Respiratory Syndrome Coronavirus 2 (SARS-CoV-2), is only the latest example of the speed with which a pathogen can travel around the globe causing successive waves of outbreaks (5).

Despite the recent emergence of this pandemic, community spread of COVID-19 is well recognized but transfusion transmission has yet to be reported (4,6). In the early days of the pandemic experts debated whether asymptomatic transmission was possible, a necessary precondition for transmission through transfusion given rigorous donor screening practices (7). That question; however, is no longer debated as the degree of community transmission in Italy and now New York have accelerated despite symptomatic screening. COVID-19 is characterized by viral shedding which starts during an initial asymptomatic phase that can last more than 14 days, followed by active disease and post-resolution viral shedding that can persist for up to 37 days (7). Furthermore, findings of viral RNA in multiple bodily fluids tested including blood raises considerable concern regarding the safety of convalescent plasma (8). In one study, viral RNA was detected in the blood of 96.8% of affected patients (9). While SARS-CoV, the causative agent of the Severe Acute Respiratory Syndrome (SARS) outbreak of 2002 and the Middle East Respiratory Syndrome Coronavirus (MERS-CoV) have not been documented to cause transfusion-transmitted disease, those pathogens resulted in higher mortality, but lower infectivity due to a lower binding efficiency with the angiotensin-converting enzyme 2 (ACE2) and dipeptidyl peptidase 4 (DPP4) receptors, respectively. By contrast, SARS-CoV-2 binding strength to the ACE2 receptor is higher, suggesting a cause for the greater propensity for human to human transmission (10,11). Observations regarding the absence of documented transfusion transmission of SARS and MERS may not be a good indication of whether COVID-19 poses a threat to the blood supply.

Pathogen reduction with riboflavin and ultraviolet light (R+UV PRT) has demonstrated excellent activity against MERS CoV, suggesting that R+UV PRT could be protective against the possibility of transfusion transmission of SARS-CoV-2 (12). In this study, we used infectivity assays to test the efficacy of R+UV PRT to reduce the level of infectious SARS-CoV-2 inoculates in plasma and whole blood.

## MATERIALS AND METHODS

### Plasma Products

Nine plasma products were collected at an accredited blood bank and shipped to Colorado State University on dry ice. The products were classified as PF24, prepared from whole blood products collected in Citrate Phosphate Double Dextrose (CP2D) and frozen within 24 hours after phlebotomy to ≤ −20°C. Products were thawed in a water bath at 36°C and pooled in sets of 3 by ABO type to create 3 unique pools. This was performed for two distinct volumes of plasma with average of 170 mLs and average of 250 mLs each.

### Whole Blood Products

Three non-leukoreduced whole blood (WB) products were collected in Citrate Phosphate Dextrose (CPD) anticoagulant at an accredited blood bank and shipped to Colorado State University at room temperature.

### SARS-CoV-2 Propagation

Virus (isolate USA-WA1/2020) was acquired through BEI Resources (product NR-52281) and amplified in Vero cell culture. Medium harvested from infected cells 3-4 days after inoculation was clarified by centrifugation, supplemented with FBS to 10% and frozen to −80°C in aliquots. All virus propagation occurred in a BSL-3 laboratory setting.

### Pathogen Reduction Process, Plasma

After pooling, each pool was divided in 3 equal volumes of 175 mL dispensed into an illumination bag (Mirasol Illumination Bag, Terumo BCT, Lakewood, CO). Riboflavin solution (35 mL, 500 μmol/L) was added to each product, followed by inoculation with 5 mL SARS-CoV-2 virus, and the bags were placed into the Illuminator (Mirasol PRT System, Terumo BCT, Lakewood, CO) for treatment with UV light. The products in each set of 3 were treated to either 30%, 60%, or 100% of the total recommended light dose, calculated based on the volume of each product (a full treatment consists of exposure to 6.24 J/mL UV light). Samples were removed from each product pre- and post-illumination for viral titer determination via plaque assay. All processing occurred in a BSL-3 laboratory setting.

### Pathogen Reduction Process, Whole Blood

Each WB product was transferred to an illumination bag per the manufacturer’s instructions. Riboflavin solution (35 mL, 500 μmol/L) was added to each product, followed by inoculation with 20 mL SARS-CoV-2 virus, and the bags were placed into the illuminator for treatment with UV light-calculated using the measured hematocrit and volume of each product - to a dose of 80 J/mL _RBC_. Samples were removed from each product pre- and post-illumination for viral titer determination via plaque assay. All processing occurred in a BSL-3 laboratory setting.

### Viral Plaque Assay

All pre- and post-illumination samples were serially diluted in sterile PBS. Plaque assays were performed using Vero cells at confluency in 6-well cell culture plates. Briefly, plates were washed with sterile PBS. All samples were then plated in duplicates at 100 µL per well. Plates were incubated at 37°C for 45 minutes with occasional rocking. Then 2 mL of 0.5% agarose in minimal essential media (MEM) containing 2% FBS and antibiotics was added per well. Plates were incubated at 37°C for 72 hours. The cells were fixed with 10% buffered formalin, followed by the removal of the overlay, and then stained with 0.2% crystal violet to visualize plaque forming units (PFU). All assays were performed in BSL-3 laboratory setting. Figure 1.

**Figure 1:**
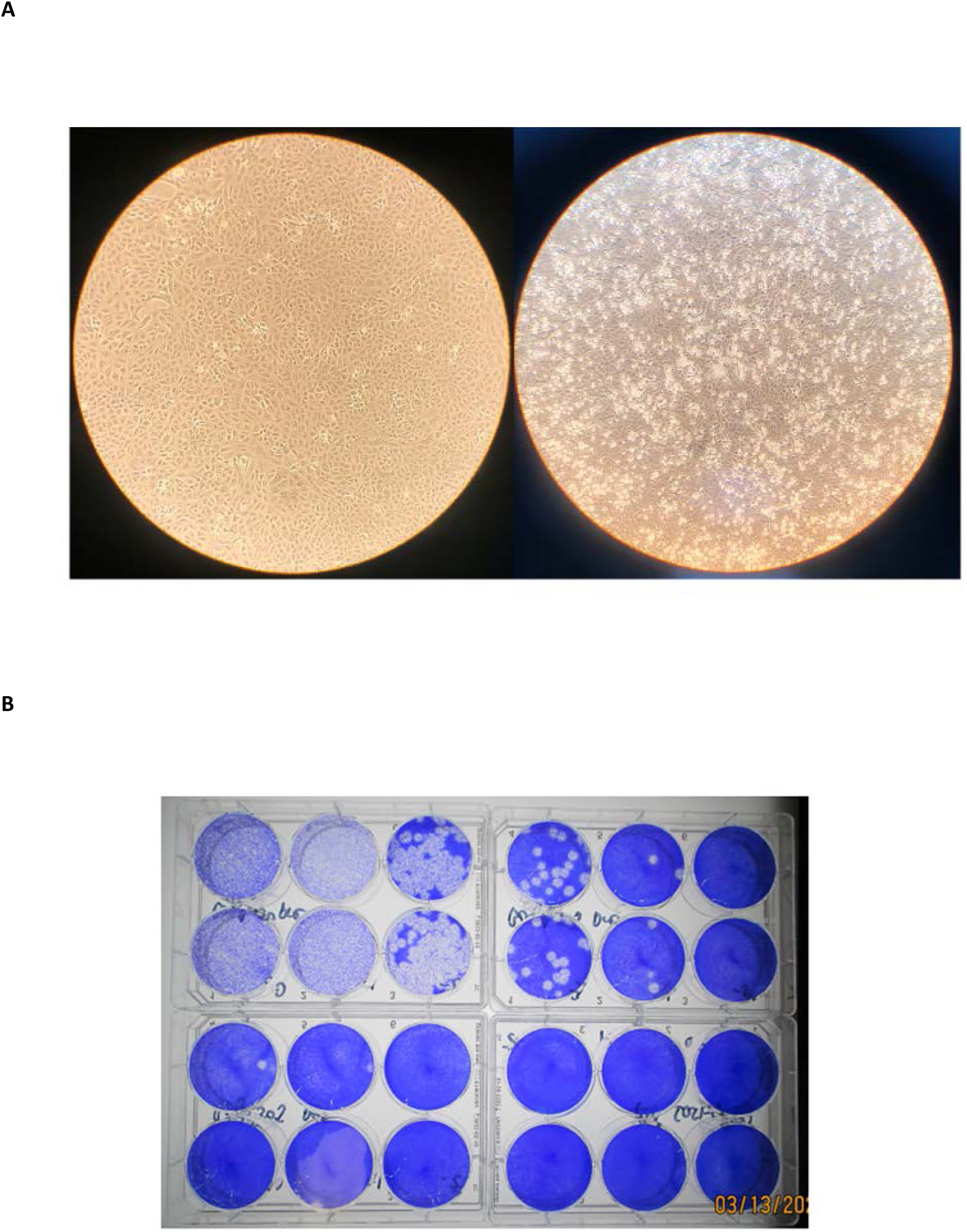
SARS-CoV-2 Cultures and Plaque Assay for Titer Determinations. A) Left-Vero cells at confluency, uninfected; Right-SARS-CoV-2 infection in Vero cells. 3 days after inoculation. CPE present. Cells at 40x magnification B) Plaque assay results from SARS-CoV-2 in media with R + UV treatment. Top left and right-virus titration, pre-pathogen reduction; Bottom left-pathogen reduction at 50J; Bottom right pathogen reduction at 100J.

### Calculation of Limit of Detection and Log Reduction

When the posttreatment samples were negative for the presence of virus, the limit of detection had been reached. All values at the limit of detection were considered less than or equal to the calculated limit of detection. The theoretical limit of detection and overall log reduction was calculated using the following equations:

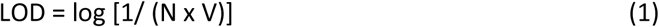

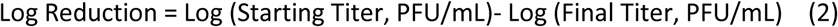

where N is the number of replicates per sample at the lowest dilution tested; V is the volume used for viral enumeration (volume inoculated/ well in mL). No cytotoxicity was observed at the zero dilution, hence all replicates for determination of LOD and final titer were done at zero dilution with 10 replicates using 0.1 mL per replicate well.

## RESULTS

Data on dilution factors for virus titration into plasma and whole blood are included in Table 1. This data indicates reasonable recovery of the stock virus titer after dilution of the virus stock into the blood products. Values between predicted and observed titers in the recovered virus from stock sample dilution into plasma and whole blood were on the order of +/-0.45 log_10_, indicating that adsorption of virus to protein or cellular components is unlikely.

**Table 1:** Stock titer values and dilution factors in blood products utilized in this study.

|  | **Plasma Study** | **Whole Blood Study** |
| --- | --- | --- |
| Stock Titer (day of)<br>(pfu/mL) | $6 \times 10^5$ | $1.5 \times 10^5$ |
| Pre-treatment Titer<br>(pfu/mL) | $5.82 \times 10^3$ | $1.83 \times 10^4$ |
| Product Volume<br>(medium + RB) (mL) | (170+35)<br>205 | (460+35)<br>495 |
| Volume Virus used<br>(mL) | 5 | 20 |
| Total Product Volume<br>(mL) | 210 | 515 |

The *in vitro* plaque assays demonstrated that pathogen reduction of plasma inoculated with SARS-CoV-2 was able to reduce infectious titers to the limit of detection by ≥ 3.77 +/-0.27 and ≥ 3.76 +/-0.15 log_10_ at 60% and 100% of the UV dose recommended by the manufacturer, respectively at volumes of 170 mLs (Table 2). The fact that the limit of detection was achieved in plasma at 60% of the recommended UV dose suggests that higher starting viral titers could have resulted in higher log kill. At higher volumes of plasma and higher spike titers, the level of reduction observed was ≥ 4.72 +/-0.24 and ≥ 4.79 +/-0.15 log_10_ at 60% and 100% of the UV dose, respectively (Table 3). Similarly, R + UV PRT was effective at reducing infectious viral loads in whole blood with a mean log reduction of 3.30+/-0.26 log_10_ in infectious titers (Table 4) but did not achieve complete inactivation, suggesting that the maximum log reduction was achieved in those samples. The level of reduction observed generally exceeded 3 log_10_ for both plasma and whole blood. Given the current limits on detection using PCR methods for screening, these results suggest at a minimum coverage for potential window period transmission risk.

**Table 2:** Log Reduction in SARS-CoV-2 coronavirus titers after pathogen reduction technology treatment of pooled plasma unit donations at volumes of 170 mLs plasma.

| **Starting Titer** | **4.63 x 10<sup>3</sup> PFU/mL** | **6.67 x 10<sup>3</sup> PFU/mL** | **6.17 x 10<sup>3</sup> PFU/mL** |
| --- | --- | --- | --- |
|  | **30% Energy Dose**<br>**Log Reduction**<br>**(N=3)** | **60% Energy Dose**<br>**Log Reduction**<br>**(N=3)** | **100% Energy Dose**<br>**Log Reduction**<br>**(N=3)** |
| **Mean** | **2.86** | **≥ 3.77** | **≥ 3.76** |
| **SD** | **0.27** | **0.27** | **0.15** |

Replicates consisted of pooled plasma units spiked with a known quantity of SARS-CoV-2 coronavirus. One bag from each pool was treated independently at each of the energy levels indicated at 30%, 60% or 100% of the full energy dose of 6.24 J/mL. Product volume at 170 mLs of plasma. Samples at limit of detection at 60% and 100% of UV dose.
PFU = plaque forming units.

**Table 3:** Log Reduction in SARS-CoV-2 coronavirus titers after pathogen reduction technology treatment of pooled plasma unit donations at volumes of 250 mLs plasma.

| **Starting Titer** | **9.12 x 10<sup>4</sup> PFU/mL** | **5.37 x 10<sup>4</sup> PFU/mL** | **6.31 x 10<sup>4</sup> PFU/mL** |
| --- | --- | --- | --- |
|  | **30% Energy Dose**<br>**Log Reduction**<br>**(N=3)** | **60% Energy Dose**<br>**Log Reduction**<br>**(N=3)** | **100% Energy Dose**<br>**Log Reduction**<br>**(N=3)** |
| **Mean** | **2.61** | **≥ 4.72** | **≥ 4.79** |
| **SD** | **0.12** | **0.24** | **0.15** |

Replicates consisted of pooled plasma units spiked with a known quantity of SARS-CoV-2 coronavirus. One bag from each pool was treated independently at each of the energy levels indicated at 30%, 60% or 100% of the full energy dose of 6.24 J/mL. Product volume at 250 mLs of plasma. Samples at limit of detection at 60% and 100% of UV dose.
PFU = plaque forming units.

**Table 4:** Log Reduction in SARS-CoV-2 coronavirus titers after pathogen reduction technology treatment of whole blood unit donations.

|  | **†Pre-treatment**<br>**PFU/mL** | **Log Reduction** |
| --- | --- | --- |
| **Product 1** | **2.50 x 10<sup>4</sup>** | **3.55** |
| **Product 2** | **1.75 x 10<sup>4</sup>** | **3.04** |
| **Product 3** | **1.25 x 10<sup>4</sup>** | **3.32** |
|  | **Mean**<br>**SD** | **3.30**<br>**0.26** |

†Products consist of whole blood units spiked with a known quantity of SARS-CoV-2 coronavirus.
PFU = plaque forming units.

## DISCUSSION

The novel COVID-19 pandemic is unprecedented in recent memory and the disease results in a severe pulmonary syndrome in a subset of patients. The potential of SARS-CoV-2 to be transmitted through transfusions is unknown. The SARS-CoV-2 virus was detected in blood at very low levels and no documented cases of transfusion transmission have been reported in the literature (13). Despite this, the World Health Organization and the US Food and Drug Administration (FDA) recommended deferring donors based on risk of SARS due to a theoretical risk of transmission (8). The current novel coronavirus pandemic exhibits higher viral loads in blood samples and far greater human-to-human spread. Early in the course of the pandemic in the United States, FDA suggested a 28-day deferral for travel-associated risk; however, with over 85 thousand cases at the time of writing and no states free of cases, this strategy is no longer tenable.

M.B. Pagano et al. described their experience with adapting to pandemic conditions in Washington State and stated that blood donations fell precipitously as the number of cases rose exponentially (14). Shortages were resolved with a combination of decreasing demand by cancelling elective surgeries, transporting blood from other states, and careful management of the supply. With a majority of the country experiencing exponential growth in cases, this may be increasingly difficult to replicate in other areas. Community spread has accelerated since the publication of the BloodWorks Northwest experience and the ability to trace the origin of each case has been lost as the pandemic has accelerated.

For a blood industry that is dependent on the use of human volunteer donors, such an event poses an existential threat to the blood supply, albeit temporary. Reduced access to donors due to restricted movement and concerns over the risk of proximity to hospitals and donor centers place severe constraints on the ability to maintain an adequate supply. With increasing prevalence of the disease and recognition of the risk of asymptomatic viral shedding come concerns that the donations may themselves serve as a vector for disease transmission. Furthermore, diagnostic testing of the blood is not feasible as the health care system struggles to address the growing number of cases needing evaluation.

In this environment, determining whether transfusion transmission in human subjects has occurred is not possible and eliminating donors with potential asymptomatic infectivity is problematic. Despite widespread cancelations of elective cases throughout the country, blood products will still be required due to injury or illness for patients who are presumed to be naïve. Additionally, FDA released an Emergency Use Authorization for compassionate use of convalescent plasma in the most critically ill patients and has signaled the willingness to consider research studies to assess the safety and efficacy in other populations. Clinical studies to assess the potential of convalescent plasma to circumvent full blown COVID-19 in exposed high-risk subjects are under consideration; however, concerns remain regarding potential inadvertent transfusion of viable virus.

The plasma fractionation industry has experience with a high risk of infection due to the large number of plasma units pooled and have developed processing methods that inactivate or reduce the levels of infectious agents in blood products. In these manufacturing settings, blood is processed to remove or inactivate pathogens and render the resulting product safe for use. Contamination with new, emerging agents is determined via direct testing or is extrapolated based on the behaviors of similar agents treated under these conditions. Methods employed include ultrafiltration, heat inactivation and solvent-detergent treatment, all of which have been found to significantly reduce pathogen contaminants when used alone or in combination in processing of donated plasma for fractionation.

Plasma fractionation techniques are not suitable for cellular components or single donation products such as Fresh Frozen Plasma (FFP) and Frozen Plasma-24 Hours (FP-24) units. Recently, however, new pathogen reduction methods have emerged for the treatment of plasma, platelets and whole blood products that can be used for treatment of single or pooled donations such as buffy coat platelets. Riboflavin is a photosensitizer that, in the presence of UV excitation, can induce changes to DNA and RNA via electron transfer chemistry and reactive oxygen species generation. These changes result in the inability of treated agents to replicate. R + UV PRT has received market approval in Europe, Africa, Asia, and South America and is in clinical development in the United States under an Investigation Device Exemption.

Our results in this study demonstrate that R + UV PRT is effective in reducing SARS-CoV-2 infectivity to the limit of detection in plasma inoculated with virus. Pathogen reduction was also effective for whole blood units, suggesting that the riboflavin and UV method can provide a measure of safety against this enveloped, positive-sense, single strand RNA virus. In a rapidly moving environment in which cases are rising by over 15,000 cases per day at the time of writing and major cities across the country struggling to contain the spread of the pandemic, security of the blood supply is in serious question. A pathogen reduction technology would provide a measure of security that would allow donations from asymptomatic adults to continue to support the critically ill requiring transfusions. Recovered patients with antibody response could provide convalescent plasma to help protect health care workers exposed to who will be needed to contain the crisis. Passive immunity could also assist high risk patients and the critically ill to clear viral loads. Previous studies have demonstrated that R + UV PRT is a potential tool to mitigate the risk of infection from convalescent plasma. An in vitro study conducted with Ebola Virus (EBOV) in serum and whole blood from non-human primates with Ebola disease demonstrated that antibody titers were maintained within protective thresholds after R+UV PRT (15).

This study has limitations. Other blood components such as platelets and red blood cells were not assessed. Plasma and whole blood from infected patients were not assessed for infectivity. The capacity of transfused blood products from an animal infected with SARS-CoV-2 to cause disease was not assessed but is the focus of ongoing studies in our labs. The limit of detection was reached in the plasma experiments thus the full range of pathogen inactivation was not measured but is greater than 4.79 ± 0.15 logs at the UV dose recommended by the manufacturer.

SARS-CoV-2 has demonstrated frequent mutations since it was first recognized in December of 2019, evolving into two different strains (designated L and S), suggesting a possible propensity to develop subtypes that could result in seasonal variability and lack of immunity to the new strains for those exposed to the previous strains (16). To date, the differences in the two strains appear to be related to infectivity and the impact on conferred immunity is not clear. In the setting of this rapidly expanding pandemic, a significant number of severely ill patients, and unclear transmissibility in blood, experts are advising caution. Even though a mere month has passed since the article by Drs. Dodd and Stramer were published, it is becoming clear that deferment is becoming increasing untenable as a strategy to address the risk (6). Pathogen reduction may be the only viable solution to protect the blood supply during this crisis.

## CONCLUSIONS

Pathogen reduction with riboflavin and UV light results in high levels of reduction of SAR-CoV-2 infectivity in inoculated plasma and whole blood. While the infectivity of blood from COVID-19 patients and its transmissibility via transfusion is not proven, the presence of viral RNA in blood is of concern and rapidly expanding community spread combined with a long asymptomatic latent period is limiting the ability of blood centers to identify who should be deferred. The rapid rise of cases suggests the US is still in the exponential phase of pandemic. Pathogen reduction could be a viable strategy to protect the blood supply during the COVID-19 pandemic.

## ACKNOWLEDGEMENTS

This work was supported by the Congressionally Designated Medical Research Program under grant number PR180446 - Indications Against Highly Pathogenic Agents for a Transportable Pathogen Reduction and Blood Safety System for Whole Blood.

We thank BEI for providing the SARS-CoV-2 utilized in these studies. The reagent was deposited by the Centers for Disease Control and Prevention and obtained through BEI Resources, NIAID, NIH: SARS-Related Coronavirus 2, Isolate USA-WA1/2020, NR-52281.

